# Hippocampus and amygdala fear memory engrams re-emerge after contextual fear relapse

**DOI:** 10.1101/2019.12.19.882795

**Authors:** Yosif Zaki, William Mau, Christine Cincotta, Emily Doucette, Stephanie L. Grella, Emily Merfeld, Nathen J. Murawski, Monika Shpokayte, Steve Ramirez

**Affiliations:** Department of Neuroscience, School of Medicine at Mount Sinai, New York, NY, 10029; Department of Psychological and Brain Sciences, Boston University, Boston, MA, 02215

## Abstract

The formation and extinction of fear memories represent two forms of learning that each engage the hippocampus and amygdala. How cell populations in these areas contribute to fear relapse, however, remains unclear. Here, we demonstrate that, in male mice, cells active during fear conditioning in the dentate gyrus of hippocampus exhibit decreased activity during extinction and are re-engaged after contextual fear relapse. *In vivo* calcium imaging reveals that relapse drives population dynamics in the basolateral amygdala to revert to a network state similar to the state present during fear conditioning. Finally, we find that optogenetic inactivation of neuronal ensembles active during fear conditioning in either the hippocampus or amygdala is sufficient to disrupt fear expression after relapse. These results suggest that fear relapse triggers a partial re-emergence of the original fear memory representation, providing new insight into the neural substrates of fear relapse.

## Introduction

The biological capacity to produce adaptive behavioral responses in actively changing environments is critical to an animal’s survival. Contextual fear conditioning (CFC) is a form of learning whereby an animal learns to associate a conditioned stimulus (i.e. a context) with an unconditioned aversive stimulus (e.g. foot shocks) to produce a conditioned response to the conditioned stimulus (e.g. freezing). Conditioned responses can be mitigated through extinction learning via repeated exposure to the conditioned context in the absence of the foot shock. However, while extinction learning can be effective at attenuating fear, animals are susceptible to fear relapse under several conditions, including exposure to stressors, the passage of time, and re-exposure to the unconditioned stimulus (Bouton & Bolles, 1979a; Travis D. Goode, Jin, & Maren, 2018; T. D. Goode & Maren, 2014; Halladay, Zelikowsky, Blair, & Fanselow, 2012; Rescorla & Heth, 1975; Vervliet, Craske, & Hermans, 2013). This observation in rodents shares numerous similarities to clinical observations: exposure therapy – a clinical analog to extinction learning – can be effective at reducing fear in subsets of patients with anxiety disorders or post-traumatic stress disorder. However, many patients are still susceptible to fear relapse following successful exposure therapy (Haaker, Golkar, Hermans, & Lonsdorf, 2014; Kearns, Ressler, Zatzick, & Rothbaum, 2012; LaBar & Phelps, 2005). Despite an extensive body of literature investigating the neural substrates of fear and extinction learning (Bocchio, Nabavi, & Capogna, 2017; LeDoux, 2000; Maren, 2001; Maren & Holmes, 2016; Milad & Quirk, 2012; Quirk & Mueller, 2008), the neural substrates underlying fear relapse are comparatively less understood.

Fear reinstatement is a form of fear relapse whereby extinguished fear of a conditioned stimulus (CS) returns following re-exposure to the unconditioned stimulus (US) alone (Rescorla & Heth, 1975). Fear reinstatement has been discussed as being a non-associative learning phenomenon driven by a changing US representation strength across conditioning and extinction (Rescorla & Cunningham, 1977). It has also separately been proposed that the CS mediates reinstatement during recall after US presentation, by retrieving the CS-US memory (Bouton, 1993; Bouton & Bolles, 1979b; Westbrook, Iordanova, McNally, Richardson, & Harris, 2002). The neural circuitry underlying fear relapse has also been studied in recent years. It has been demonstrated, for example, that pharmacological activation of noradrenergic activity is sufficient to drive fear relapse after extinction (Giustino, Fitzgerald, Ressler, & Maren, 2019), that dopamine-1 receptor blockade in the infralimbic cortex prevents fear reinstatement (Hitora-Imamura et al., 2015). and that activity of the bed nucleus of the stria terminalis is necessary for fear reinstatement (T. D. Goode, Kim, & Maren, 2015). However, how brain regions central to emotional memory processing – such as the amygdala and hippocampus – contribute to fear relapse has been understudied. Nonetheless, it has been shown that pharmacological inactivation of either the prelimbic cortex, dorsal CA1, or ventral hippocampus disrupts fear relapse (Fu et al., 2016), and blockade of either mRNA or protein synthesis in the CA1 of the hippocampus prevents fear relapse (Cammarota, Bevilaqua, Kerr, Medina, & Izquierdo, 2003). More broadly, similar sets of brain regions, including the amygdala and hippocampus, are involved in the relapse of both fear and drug intake (T. D. Goode & Maren, 2019). However, whether and how discrete neuronal populations active during fear conditioning contribute to fear reinstatement remains incompletely understood.

Previous studies have demonstrated that cells in the dorsal dentate gyrus of the hippocampus (DG), in the CA1 of the hippocampus, and in the basolateral amygdala (BLA) that are active during fear conditioning (hereafter referred to as the DG, CA1, and BLA fear ensembles) are preferentially re-activated during fear memory recall (Ghandour et al., 2019; Nakazawa, Pevzner, Tanaka, & Wiltgen, 2016; Ramirez et al., 2013; Reijmers, Perkins, Matsuo, & Mayford, 2007), and are necessary and sufficient for the expression of defensive behaviors such as freezing (Denny et al., 2014; Ohkawa et al., 2015; Redondo et al., 2014). Additionally, recent evidence has indicated that extinction learning is mediated by interactions between local BLA interneurons and a BLA fear ensemble (Davis, Zaki, Maguire, & Reijmers, 2017), while a new set of cells simultaneously emerges in both the hippocampus (Khalaf et al., 2018; Lacagnina et al., 2019; Tronson et al., 2009) and BLA (Herry et al., 2008), possibly to encode extinction learning. It has also been shown that, after fear extinction, activation of the brain-wide neuronal ensemble active during fear conditioning is capable of driving freezing, suggesting that there is a latent representation of the fear memory that can be activated exogenously after extinction (Yoshii, Hosokawa, & Matsuo, 2017). Whether fear reinstatement re-engages the original memory-encoding neuronal population or gives rise to a new representation, however, remains unclear.

## Results

### Hippocampal Cells Active During Fear Conditioning Are Less Active After Fear Extinction And Re-Engaged After Fear Reinstatement

We first developed a five-day behavioral protocol for fear reinstatement, a model of fear relapse in rodents (Rescorla & Heth, 1975). Mice underwent CFC on day 1, followed by two subsequent extinction (EXT) sessions on days 2 and 3. 24 hours after the second EXT session, mice received an immediate shock (IS) in a novel context to reinstate the original fear memory. The following day, mice were placed in a post-reinstatement recall test (IS-Recall) to measure the return of fear (Figure 1a, bottom behavioral schedule; see Methods). Reinstatement led to an increase in freezing in the original conditioned context (Supplementary Figure 1a-e) and mice froze more in the original conditioned context over a novel context after reinstatement (Supplementary Figure 2a,b).

**Figure 1.**
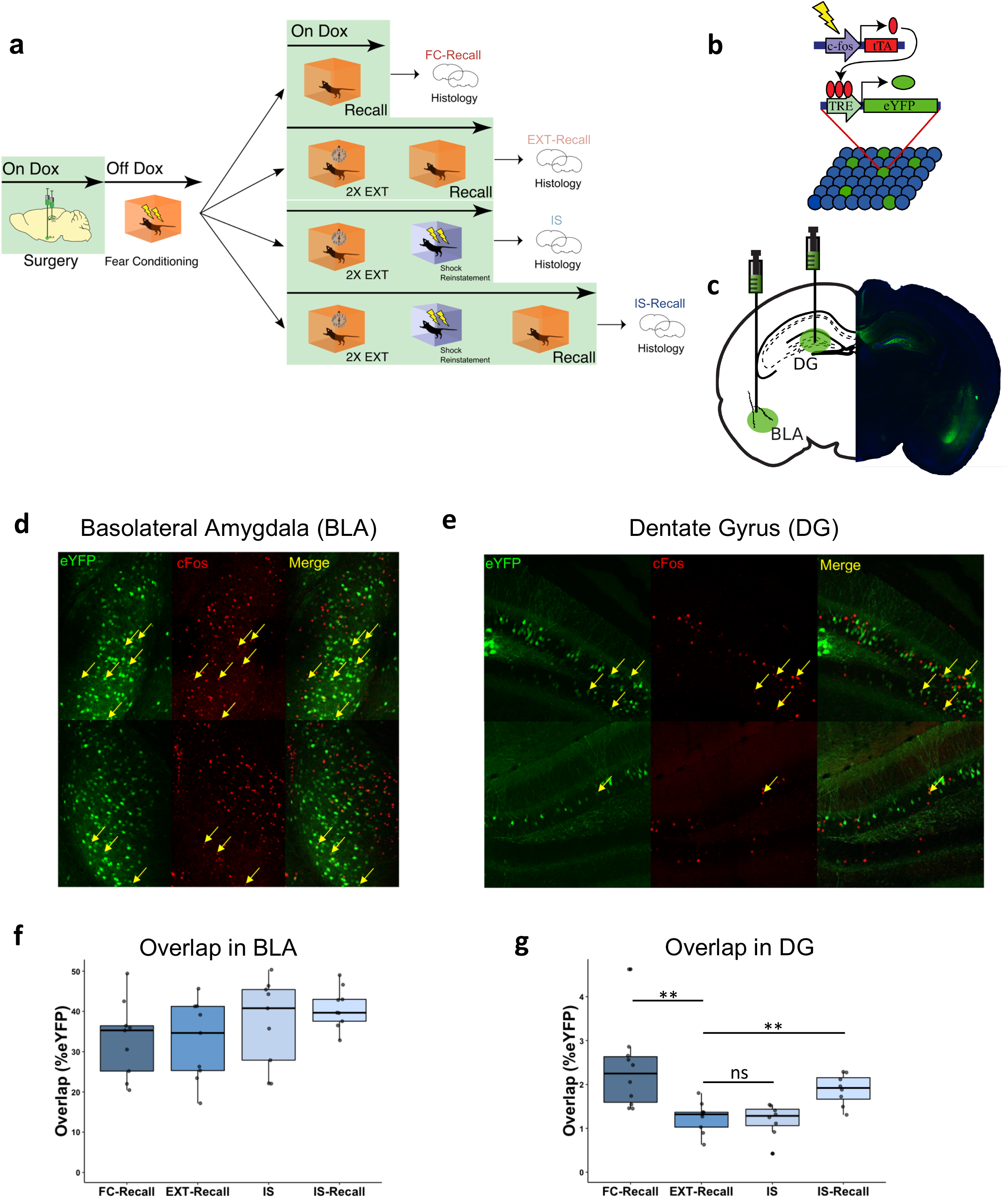
Histological characterization of fear reinstatement schedule. **(a)** Behavioral design for fear reinstatement. Mice underwent fear conditioning, and were then sacrificed at different points in the behavioral schedule and had tissue stained for c-Fos. **(b)** Schematic of viral strategy. A viral cocktail of AAV9-c-Fos-tTA and AAV9-TRE-eYFP was infused into the DG and BLA for activity-dependent induction of eYFP. **(c)** Representative microscope image for the injection sites. **(d)** Example confocal images of BLA sections. Images from left to right: virus-labeled cells (eYFP), c-Fos^+^ cells (cFos), merged green and red channels (Merge). Yellow arrows designate double-positive cells. Top images are representative of high overlap (from FC-Recall group), and bottom are representative of low overlap (from EXT-Recall group). **(e)** Same as (**d)** but for DG sections. **(f)** Quantitative analysis of overlap between FC-tagged BLA cells and c-Fos^+^ BLA cells in each group. The amount of overlap between FC-tagged (eYFP+) cells and c-Fos^+^ cells remained unchanged across all behavioral groups. (n = 8-10 mice per group; F_3,32_ =1.607; one-way ANOVA). Counts were calculated as % of eYFP^+^ cells that were also c-Fos^+^. Each data point represents one mouse. **(g)** Same as quantification in (**f)**, but for DG. Overlap between FC-tagged cells and c-Fos^+^ cells was high during recall after fear conditioning (FC-Recall) and significantly decreased following EXT (EXT-Recall). While overlap remained low during the reinstating shock (IS), it significantly increased during Recall after reinstatement (IS-Recall). (n = 8-10 mice per group; F_3,31_ = 7.703, p = 0.0006; one-way ANOVA followed by pairwise t-tests corrected with Benjamini-Hochberg method to correct for multiple comparisons; **P < 0.01, ns P > 0.05).

Next, we determined if the cells active during fear conditioning were preferentially reactivated after mice underwent extinction and subsequent reinstatement. To that end, we tagged cells active during fear conditioning by injecting an activity-dependent viral cocktail of AAV9-c-Fos-tTA and AAV9-TRE-eYFP in the DG and BLA of adult male mice (Figure 1b,c). This virus enabled expression of eYFP in cells sufficiently active to express the immediate early gene c-Fos, which is under the repressive control of the antibiotic doxycycline (DOX)(Reijmers et al., 2007). We then measured immunoreactive c-Fos and calculated overlap between the set of cells active during CFC (eYFP-expressing cells) and cells active during different stages of the behavioral schedule (c-Fos-expressing cells) (Figure 1d,e).

Previous reports have shown that the number of BLA cells active during both fear conditioning and fear memory recall correlates with freezing levels (Reijmers et al., 2007). Thus, we reasoned that if reinstatement re-engages the fear ensemble, the set of cells active during fear conditioning would be active again following reinstatement. Surprisingly, we found that cells active during CFC were no differently reactivated throughout the behavioral schedule (Figure 1f). In the DG, however, we observed significant overlap between the cells active during CFC and those active during fear memory recall, as previously reported (Ramirez et al., 2013) (Figure 1g). In support of the notion that the dorsal DG processes changes in environmental contingencies (Fanselow & Dong, 2010), this overlap substantially decreased after EXT. While overlap remained low after IS, it significantly increased when mice were given the IS and were placed back into the original conditioned context the following day, suggesting that fear reinstatement may re-engage the set of cells originally active during fear conditioning in the DG (Figure 1g). A similar analysis where the number of overlapping cells was normalized to the area of interest (rather than normalized to the number of FC-tagged cells) yielded similar results (Supplementary Figure 1f,g). Overall expression of eYFP and cFos was stable across groups in the DG, and while eYFP expression was stable across groups in the BLA, mice had significantly higher levels of cFos during post-reinstatement recall (Supplementary Figure 1h-k).

### Population Activity During Fear Conditioning Correlates With Activity After Fear Reinstatement But Not After Extinction Learning

Whereas our c-Fos-based labeling system allowed comparisons between activity of cells across two discrete timepoints with high spatial resolution, it was incapable of measuring activity at finer timescales due to the slow kinetics of immediate-early gene expression relative to real-time neural activity. To overcome this weakness, we next utilized an *in vivo* calcium (Ca^2+^) imaging approach to record real-time neuronal activity in the BLA or CA1 in freely moving mice during exposures to both a conditioned context and a neutral context where no shocks were delivered (Figure 2a-d, Supplemental Table 1). We tracked these cells longitudinally over the course of the reinstatement schedule in order to determine whether similar activity patterns were expressed during fear conditioning and reinstatement (Sheintuch et al., 2017) (Supplementary Figure 4). There were no differences in overlaps of BLA cells detected with calcium imaging for each session pair (Supplementary Figure 4c, f), consistent with our cFos immunostaining results (Figure 1f). However, the moment-by-moment activity patterns of the recorded neuronal populations might correlate with an initial population state reflecting fear acquisition during CFC. To define this initial population state, we constructed Ca^2+^ transient rate population vectors from the CFC session for each mouse. Then, to compare extinction and post-reinstatement recall states to CFC, we correlated population vectors from EXT and Recall (in 30 s non-overlapping time windows) to the CFC population vector. We found that over EXT, the population states in the BLA and CA1 followed a trajectory that gradually deviated from its CFC state, supporting the idea of a network-wide transformation over extinction (Grewe et al., 2017; Hartley et al., 2019; Herry et al., 2008; Tronson et al., 2009) (Figure 2e, left). However, during Recall, the BLA population in each mouse rebounded towards the CFC network state to an extent greater than expected by chance (Figure 2e, right). This rebound effect was absent in a neutral context (Figure 2g-h), demonstrating that the conditioned context drove these dynamics primarily in BLA, and in CA1 of some mice (Figure 2f, right). Additionally, a regression continuity analysis (Thistlethwaite & Campbell, 1960) (see Methods) detected a significant change point in activity patterns at the EXT2-Recall border, suggesting that the network drastically shifted to a different state during that transition (Figure 2e-f, left). No such divergence was detected across EXT1 to EXT2 in BLA, indicating a relatively steady transition across the two EXT sessions.

**Figure 2.**
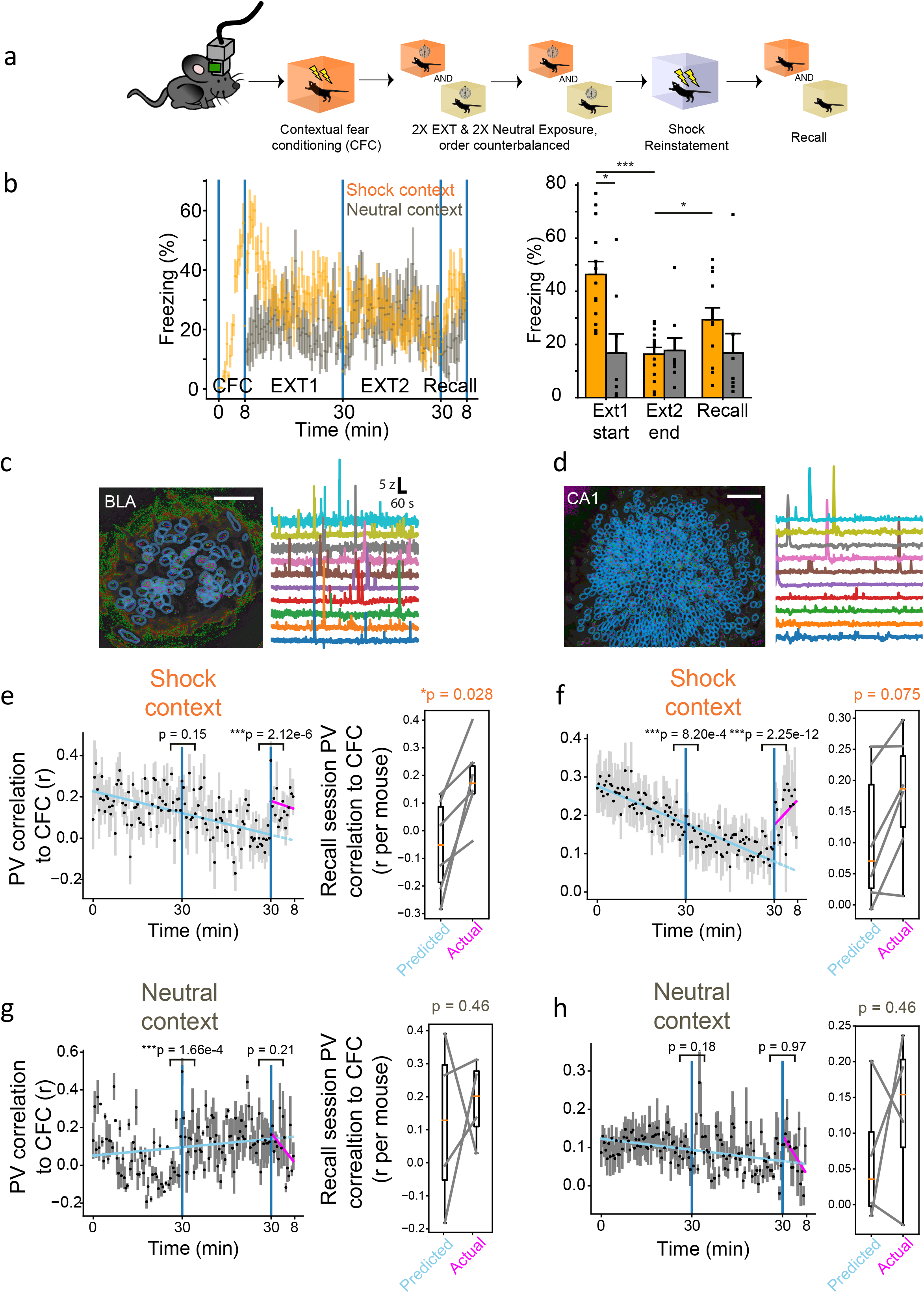
BLA activity patterns change over extinction but resemble the fear conditioning state during fear relapse. **(a)** Behavioral schedule for Ca^2+^ imaging cohort. **(b)** Left, freezing time course of fear reinstatement paradigm (*n* = 12 mice). Right, pooled freezing. EXT1 vs. EXT2, Wilcoxon signed-rank test, P = 0.0015; EXT1 vs neutral EXT1, P = 0.017; EXT2 vs. Recall, P = 0.012. Data are means + standard error of the mean. **(c)** Left, example field of view in BLA-implanted mouse, depicted as maximum projection of CFC imaging session. Blue outlines indicate cell masks. Scale bar = 100 microns. Right, fluorescence traces of 10 example cells. **(d)** Same as (**b**), but for CA1. **(e)** Left, Pearson correlation coefficients between population vectors during CFC and population vectors during EXT and Recall (*n* = 6 BLA mice). Each data point represents a measure of population similarity to CFC (30 s time bins). Two piecewise regressions were fit each to EXT and Recall. P-values above indicate regression discontinuity analysis results on discontinuities between population vector correlations across sessions (EXT1 to EXT2 and EXT2 to Recall). Right, box plots of population vector similarity (predicted and empirical). Each data point represents a mouse’s predicted r value (from the EXT regression) for Recall compared to the actual observed r value during Recall (Wilcoxon signed-rank test, P = 0.028). **(f)** Same as (**e**), but for CA1. **(g)** Same as (**e**), but for the neutral context. **(h)** Same as (**f**), but for the neutral context.

Using an algorithm for extracting co-active neurons from simultaneously recorded cells (Lopes-dos-Santos, Ribeiro, & Tort, 2013), we also identified neuronal ensembles during individual sessions prior to Recall (CFC, EXT1, and EXT2). We hypothesized that the ensembles identified during CFC might predict freezing behavior during Recall, while ensembles identified during late extinction sessions would not. On the Recall session, we correlated the activity of these ensembles to freezing and found that the activity of ensembles extracted during CFC and EXT1 reliably predicted relapse freezing, but later ensembles extracted during EXT2 did not (Supplementary Figure 3c). No CA1 ensembles from any session predicted freezing nor did BLA or CA1 ensembles predict freezing in the neutral context (Supplementary Figure 3d). This suggests that BLA activity patterns contributed to expression of fear relapse, but only before extinction training modified these patterns. Overall, these data indicated that context-specific reinstated fear was associated with the emergence of network states in the BLA that resembled network states during fear conditioning, suggesting that a relapsed fear memory may be represented by a similar trace as the original fear memory.

### Optogenetic Inhibition of the BLA DG, or CA1 Fear Ensemble Disrupts Freezing After Fear Reinstatement

Finally, we sought to determine whether the activity of cells active during fear conditioning was necessary for expression of reinstated fear. To do this, we bilaterally injected mice in either the BLA, DG, or CA1 with a virus cocktail of AAV9-c-Fos-tTA and AAV9-TRE-ArchT-eYFP to drive expression of the light-sensitive protein archaerhodopsin (ArchT) in cells active during CFC, and subsequently implanted optic fibers above the injection sites (Figure 3a,b). Mice then underwent two EXT sessions, the reinstating shock, and recall the following day (Figure 3c). Mice in all three experimental groups (i.e. BLA, DG, CA1) showed significant suppression of freezing during optical inhibition (Figure 3d,f,h). In the BLA and DG, this manipulation was reversible, as freezing increased again in the following light-off epoch (Figure 3d,e). In the CA1, however, freezing did not increase again once the light stimulation ended (Figure 3h). eYFP controls did not show this decrease in freezing during optical inhibition, confirming that the behavioral effect was dependent on expression of ArchT (Figure 3e,g,i). Difference scores comparing freezing during light on vs light off epochs confirmed that mice which received inhibition of the fear ensemble in either the BLA, DG, or CA1 froze significantly less during light-on period compared to light-off periods, and control groups froze no differently across the two light conditions (Figure 3j).

**Figure 3.**
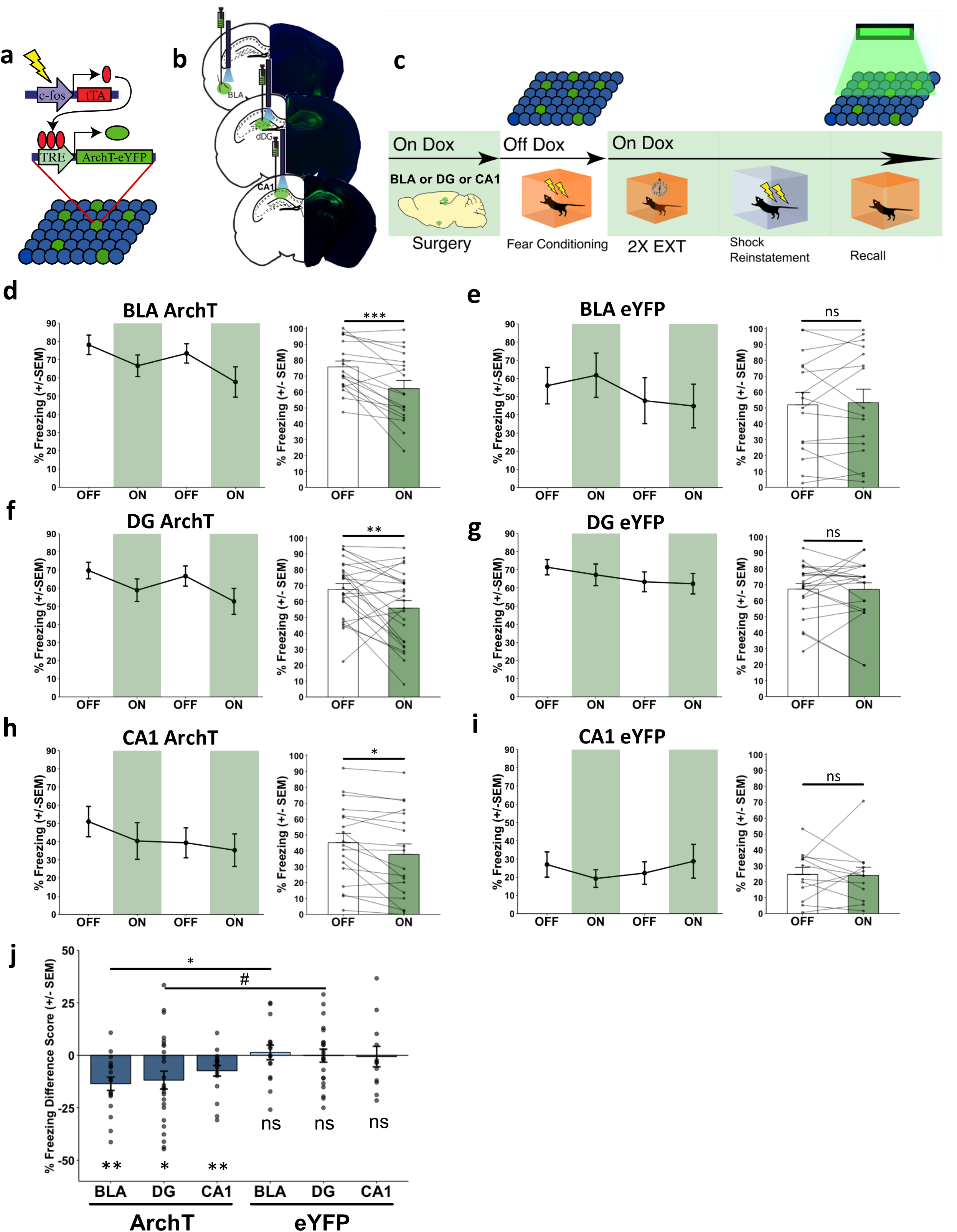
Optical inhibition of the DG or BLA fear ensemble disrupts reinstated fear. **(a)** Schematic of viral strategy. A virus cocktail of AAV9-c-Fos-tTA and AAV9-TRE-ArchT-eYFP was infused into either the DG or BLA for activity-dependent expression of ArchT-eYFP. **(b)** Representative microscope images of injection sites for the DG and BLA groups of mice. **(c)** Reinstatement behavioral schedule. Mice had the fear ensemble labeled in either the DG or BLA and inhibited during Recall after Shock Reinstatement. **(d-i)** Line graphs: 2-minute light OFF and ON epochs during Recall for the three experimental ArchT groups (BLA Exp, DG Exp, CA1 Exp) and the three no-opsin control groups (BLA eYFP, DG eYFP, CA1 eYFP). Bar graphs: Quantification of average freezing between light OFF (white bars) vs. light ON (green bars) epochs for each group. **(d)** BLA Exp Recall: mice froze significantly less during ON epochs than OFF epochs (Main effect of Light: F_(1,8)_ = 38.71, ***P = 0.0002; two-way repeated measures ANOVA of Light (OFF vs ON) & Epoch (1 vs 2); n = 9 mice). **(e)** BLA eYFP Recall: mice froze no differently during ON epochs than OFF epochs (No main effect of Light: F_(1,7)_ = 0.18, n.s., P = 0.6857; two-way repeated measures ANOVA of Light (OFF vs ON) & Epoch (1 vs 2); n = 8 mice). **(f)** DG Exp Recall: mice froze significantly less during ON epochs than OFF epochs (Main effect of Light: F_(1,11)_ = 10.16, **P = 0.0086; two-way repeated measures ANOVA of Light (OFF vs ON) & Epoch (1 vs 2); n = 12 mice). **(g)** DG eYFP Recall: mice froze no differently during ON epochs than OFF epochs (No main effect of Light: F_(1,10)_ = 0.88, n.s., P = 0.3710; two-way repeated measures ANOVA of Light (OFF vs ON) & Epoch (1 vs 2); n = 11 mice). **(h)** CA1 Exp Recall: mice froze significantly less during ON epochs than OFF epochs (Main effect of Light: F_(1,8)_ = 6.46, *P = 0.0346; main effect of Epoch: F_(1,8)_ = 11.00, *P=0.0106; two-way repeated measures ANOVA of Light (OFF vs ON) & Epoch (1 vs 2); n = 9 mice). **(i)** CA1 eYFP Recall: mice froze no differently during ON epochs than OFF epochs (No main effect of Light: F_(1,5)_ = 0.03, n.s., P = 0.8688; two-way repeated measures ANOVA of Light (OFF vs ON) & Epoch (1 vs 2); n = 6 mice). **(j)** Summary graph of freezing difference scores across all groups in Figure 3. While mice in all three experimental groups (BLA, DG, CA1; dark blue bars) show significantly less freezing during light ON epochs, all eYFP control groups (BLA, DG, CA1; light blue bars) show no difference in freezing between light ON and light OFF epochs (from left to right: n = 18 scores from 9 mice, 25 scores from 13 mice, 18 scores from 9 mice, 16 scores from 8 mice, 22 scores from 12 mice, 12 scores from 6 mice; BLA Exp, t_17_ = -4.277, ***P = 0.0005; DG Exp, t_24_ = -2.781, *P = 0.0104; CA1 Exp, t_17_ = -2.9033, **P = 0.0099; BLA eYFP, t_15_ = 0.3915, n.s., P = 0.7010; DG eYFP, t_21_ = -0.05076, n.s., P = 0.9600; CA1 eYFP, t_11_ = -0.1253, n.s., P = 0.9026; one-sample t-tests, µ_0_ = 0). The BLA Exp group showed a significantly more negative difference score than the BLA eYFP group, and the DG Exp group showed a modestly more negative difference score than the DG eYFP group (three pairwise t-tests, corrected with FDR correction; BLA Exp vs BLA eYFP, DG Exp vs DG eYFP, CA1 Exp vs CA1 eYFP). BLA: t_32_ = -3.1676, *P = 0.0101; DG: t_45_ = -2.1679, #P = 0.0532; CA1: t_28_ = -1.3427, n.s., P = 0.1902.

Since the BLA is widely acknowledged as a necessary hub for fear learning (Bocchio et al., 2017), we next probed whether activity of the BLA fear ensemble during the reinstating shock was necessary or sufficient to induce fear reinstatement. To test necessity, we adopted a similar approach as above in order to express ArchT selectively within the BLA fear ensemble, and then implanted optic fibers bilaterally above BLA (Supplementary Figure 5a,b). Mice underwent FC and EXT, had the BLA fear ensemble inhibited during the reinstating shock, and were then returned to the original conditioned context to assess whether reinstatement could be prevented (Supplementary Figure 5c,d). Mice that had the BLA fear ensemble inhibited did not freeze any less during post-reinstatement recall than eYFP controls (Supplementary Figure 5e), consistent with a prior report that BLA inactivation does not prevent fear reinstatement (Laurent & Westbrook, 2010). To test sufficiency, we selectively expressed ChR2 in either the BLA fear ensemble or DG fear ensemble in separate groups of mice. Mice underwent FC and EXT, were then placed in a novel chamber, and rather than receiving the reinstating shock, mice had either the BLA or DG fear ensemble stimulated for 60 seconds. The next day, they were placed back in the original conditioned context to assess whether the stimulation could mimic reinstatement (Supplementary Figure 6a-d). Mice that had the BLA fear ensemble stimulated froze no more than eYFP controls (Supplementary Figure 6e), while mice that had the DG fear ensemble stimulated showed modest, non-significant increases in freezing relative to the eYFP controls (Supplementary Figure 6f). These results indicated that despite a crucial role for the BLA and DG fear ensembles in fear learning, activity of these populations was not sufficient to drive fear reinstatement, and activity of the BLA population was not necessary to drive reinstatement either.

To test whether the functional role for these cells emerged only after reinstatement or if inhibition of the fear ensemble could suppress freezing during extinction, we inhibited the DG or BLA fear ensemble during an extinction recall session—when low levels of freezing were still present—and observed that inhibition of the DG fear ensemble led to a mild, non-significant reduction in freezing (Supplementary Figure 7d,e,g), while inhibition of the BLA fear ensemble did not disrupt freezing (Supplementary Figure 7f). These results suggested that extinction differentially modified the BLA and DG fear ensembles, such that BLA ensemble inhibition did not disrupt freezing during extinction, while DG ensemble activity may have been actively involved in contextual fear expression following partial extinction.

## Discussion

The dynamic nature of fear memory expression constitutes a difficult problem for mitigating fear in the clinic: patients with fear-related disorders who have undergone successful treatment are still prone to relapse, and the underlying causal mechanisms facilitating fear reinstatement are largely unknown. A commonly held view is that fear extinction is not an unlearning of the original trauma; rather, a second memory develops that suppresses the original aversive memory. This raises an important notion about the nature of the ensemble regulating fear expression post-reinstatement. One idea is that the original ensemble driving fear expression and a new ensemble driving fear suppression actively compete to influence behavioral output. Under this framework, fear relapse could be the result of the fear ensemble dominating. Alternatively, fear relapse might be driven by recruitment of a new, discrete cellular population that does not involve the original fear ensemble. A likely scenario is a mixture of the two, where fear relapse materializes from a partial re-emergence of the original ensemble in parallel with recruitment of new neuronal connections (Clem & Schiller, 2016), which our c-Fos labeling, Ca^2+^ imaging, and optogenetics evidence collectively support.

Ca^2+^ imaging and c-Fos labeling during the fear reinstatement schedule enabled us to capture network dynamics from the hippocampus and amygdala over multiple timescales, shedding light on the activity of these regions over fear reinstatement. Consistent with prior reports of BLA cell populations up- and down-regulating their activity during extinction learning (Grewe et al., 2017; Herry et al., 2008), we observed decorrelation of the BLA population vector from the initial fear-encoding state over repeated exposures to the conditioned context. This time-dependent transformation has previously been depicted in numerous brain regions as “representational drift” (Driscoll, Pettit, Minderer, Chettih, & Harvey, 2017; Mankin et al., 2012; Mau, Hasselmo, & Cai, 2020; Mau et al., 2018; Rubin, Geva, Sheintuch, & Ziv, 2015; Rule, O’Leary, & Harvey, 2019), but these studies all described population states that monotonically drifted *away* from a reference session. In the present study, the BLA representation exhibited similar dynamics, but in contrast to past work, regressed its neural trajectory *back towards* the initial representation after fear reinstatement. This leads us to believe that fear reinstatement may be restoring a remote memory trace similar to how optogenetic activation can artificially induce memory retrieval (Liu et al., 2012; Ramirez et al., 2013; Redondo et al., 2014).

Interestingly, the neural patterns associated with fear expression are still retrievable after putative circuit remodeling over extinction learning and fear reinstatement (Bocchio et al., 2017; Davis et al., 2017; Hartley et al., 2019; Maren, 2015). Our ability to observe and manipulate the original fear ensemble after reinstatement is suggestive of a latent representation of the original memory that persists and coexists with the newly formed extinction memory (Lacagnina et al., 2019; Maren, 2011). However, this new extinction memory may also facilitate local synaptic remodeling that modifies the original fear memory, which may explain why inhibition of the fear ensemble in DG and BLA after extinction did not fully eliminate freezing (Supplementary Figure 7c-f) (Davis et al., 2017; Trouche, Sasaki, Tu, & Reijmers, 2013). The re-emergence of the original fear memory may also depend on strict endogenous plasticity mechanisms, which can explain why we failed to optically induce relapse through broad stimulation of the BLA fear ensemble after extinction (Supplementary Figure 6). These results suggest that the natural endogenous fear reinstatement process might require certain temporal activity patterns for modifying the original fear ensemble that could not be artificially produced through blanketed BLA stimulation alone. For example, recent work showed that sequential activity from triplets of BLA neurons preceded fear learning (Reitich-Stolero & Paz, 2019) and similar patterns may be required for fear reinstatement. Fear reinstatement may then be recruiting a subset of the original fear ensemble while forming new synaptic linkages with a novel cell population. Others have shown that unique memories reside in patterns of connectivity between memory-encoding – or engram – cells in the hippocampal system (Abdou et al., 2018; Ryan, Roy, Pignatelli, Arons, & Tonegawa, 2015; Tonegawa, Morrissey, & Kitamura, 2018). In the context of our study, reinstatement could be modifying these functional linkages to engage a new set of engram cells, possibly those that are highly excitable at the time of the experience (Cai et al., 2016; Rashid et al., 2016; Yiu et al., 2014), forming a reinstatement ensemble that is similar, but not identical, to the original fear ensemble. In accordance with this idea, post-reinstatement recall activates a large proportion, but not all, of the original fear ensemble (Figure 1f,g). Notably, past studies that have promoted learning via artificial stimulation of neural ensembles have promoted novel associations between conditioned and unconditioned stimuli (Ohkawa et al., 2015; Ramirez et al., 2013; Vetere et al., 2019). In our case, however, fear ensemble manipulation during reinstatement was meant to re-activate the latent fear memory representation (Supplementary Figures 5 & 6). Since the fear ensemble stimulation alone was insufficient to drive the re-emergence of context-specific fear (Supplementary Figure 6), and fear ensemble inhibition was insufficient to prevent reinstatement (Supplementary Figure 5), this may suggest that fear reinstatement is mediated by an alternate mechanism that requires the presentation of the unconditioned stimulus. It is possible, for example, that the nociceptive sensory inputs experienced during the immediate shock provide the necessary input to drive fear relapse (Vetere et al., 2019), after which fear expression is mediated, in part, by the original fear-encoding ensemble (Figure 3). Future studies investigating the role that sensory stimuli play for fear relapse would provide insight into this possibility.

While our optogenetic experiments suggest a re-emergence of the fear ensemble in the DG, we were unable to perform calcium imaging in DG due to the technical limitations of accessing this region without significant damage to overlying hippocampal subareas. Nonetheless, we report that CA1 exhibits only marginally significant (p = 0.075, Figure 2f) reversion of calcium dynamics to a fear conditioning-like state after reinstatement. This may be due to the functional differences between DG and CA1, with CA1 possibly responding to more contextual features of the experience (Tanaka et al., 2018) or being overall more subject to plasticity than DG. As others have found, day-to-day dynamics of spatial activity in DG are more stable than in CA1 (Hainmueller & Bartos, 2018). Until methods are developed that allow imaging of DG while still preserving superficial hippocampal areas, it remains unknown whether the real-time network activity of DG exhibit relapse-induced reactivation of fear ensembles. However, recent work agrees with our prediction that extinction suppresses fear-related activity in the DG while those activity patterns are retrieved during spontaneous recovery, another form of fear relapse (Lacagnina et al., 2019).

Further work exploring the competing interactions of cellular networks across fear learning and fear suppression could provide important insight into how the brain competes for the expression of fear throughout fear extinction and relapse. Moreover, a deeper understanding of how fear memories are modified by time and experience may help guide development of treatments for trauma-related disorders, and these findings point to hippocampal- and BLA-mediated engrams as key nodes contributing to the re-emergence of a contextual fear memory.

## Acknowledgements

We thank Dr. Joshua Sanes and his lab at the Center for Brain Science, Harvard University, for providing laboratory space within which the initial experiments were conducted, the Center for Brain Science Neuroengineering core for providing technical support, and the Society of Fellows at Harvard University for their support. We also thank Dr. Susumu Tonegawa and his lab for providing the activity-dependent virus cocktail, Dr. Chris MacDonald for consultation on behavioral schedules, and Drs. Leon Reijmers and Patrick Davis, for their help with formulating this project and for their feedback throughout. We also thank Vardhan Dani and Inscopix for their technical assistance as well as Helen Fawcett and the NSF Neurophotonics Research Traineeship Program for funding and support. This work was supported by an NIH Early Independence Award (DP5 OD023106-01), an NIH Transformative R01 Award, a Young Investigator Grant from the Brain and Behavior Research Foundation, a Ludwig Family Foundation grant, and the McKnight Foundation Memory and Cognitive Disorders award.

## Author Contributions

Y.Z. and S.R. conceived the study. Y.Z., W.M., and S.R. designed experiments and analyzed data. Y.Z., W.M., C.C., A.H., E.D., S.L.G., E.M., N.J.M., and M.S. conducted experiments. Y.Z., W.M., and S.R. wrote the manuscript; all authors edited and commented on the manuscript.

## Figure Legends

**Supplementary Figure 1.**
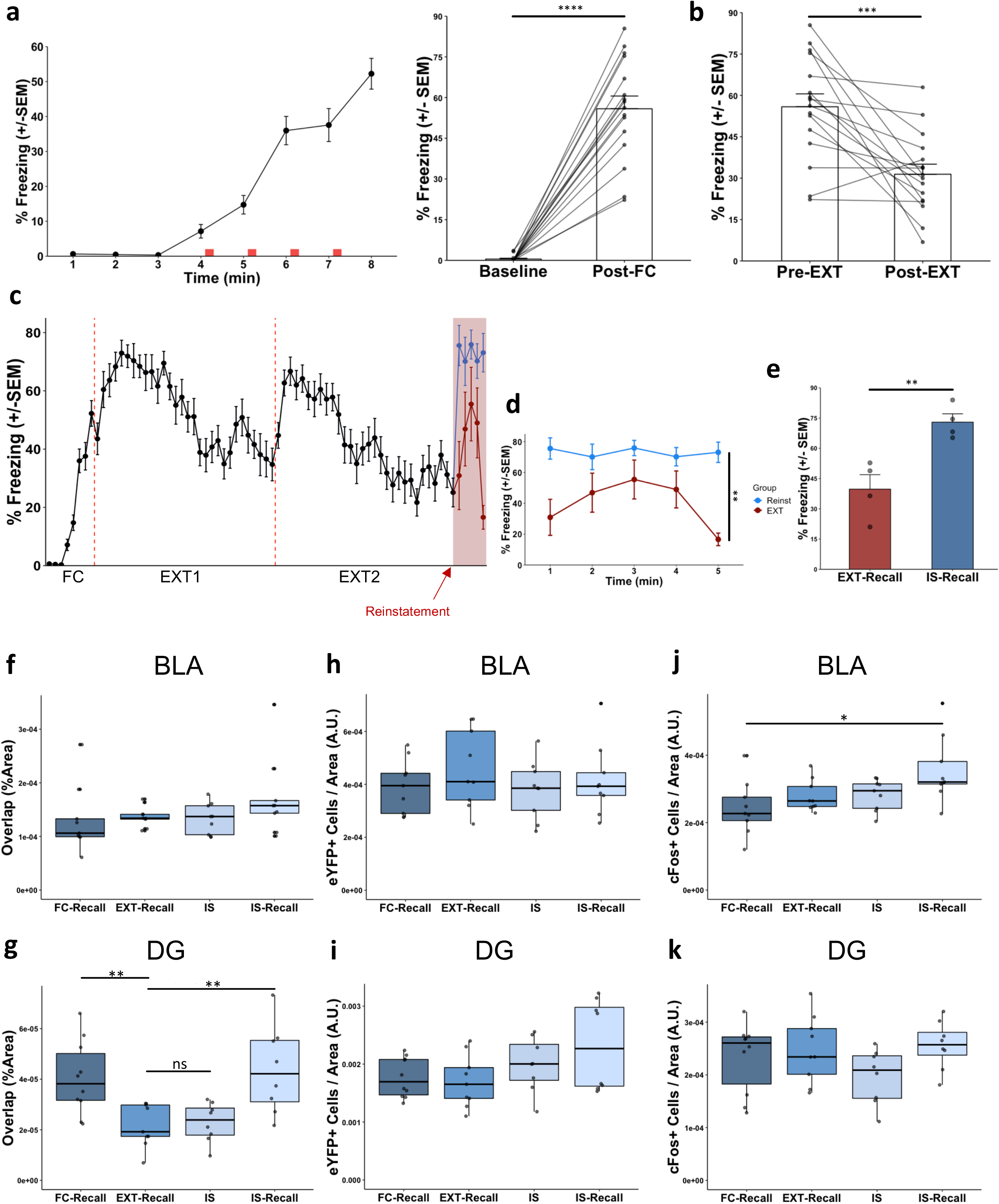
Behavior in reinstatement paradigm. **(a)** Mice underwent an 8-minute fear conditioning session (FC), with four 1.5mA shocks spaced 80-seconds apart. Compared to the first 3 minutes of FC (baseline freezing), mice froze significantly more in the first three minutes of the EXT1 session (t_15_= 11.858, ****P < 0.0001; paired t-test; n = 16 mice). **(b)** Mice underwent two 30-minute extinction sessions (EXT1 and EXT2) spaced 24 hours apart. Mice spent significantly less time freezing in the last three minutes of EXT2 as compared to the first three minutes of EXT1 (t_15_ = 4.1631, ***P = 0.0008; paired t-test; n = 16 mice). **(c)** Following EXT, one group of mice was returned to the conditioned context for Recall (EXT-Recall), while another group received an immediate shock in a novel context and was removed 60-seconds later. After 24 hours, those mice were tested in the original conditioned context for reinstatement (IS-Recall). Each data point represents mean freezing across 1-min periods. **(d)** Compared to mice that did not receive the reinstating shock, those that did showed significantly more freezing across a 5-minute Recall session (t_8_= 4.631, **P = 0.0017; unpaired t-test; n = 5 minutes for each group). **(e)** On average, mice that received the reinstating shock froze significantly more during Recall than did mice that did not receive the reinstating shock (t_6_ = 4.018, **P = 0.0070; unpaired t-test; n = 4 mice per group). **(f)** Quantification of overlap between FC-tagged (eYFP+) cells and cFos+ cells in the BLA, normalized to the area of the region of interest. As in Figure 1, the number of overlapping cells remained unchanged across behavioral conditions (n = 8-10 mice per group; F_3,32_ = 1.256, p = 0.306; one-way ANOVA). Each data point represents one animal. **(g)** Same as **(f)** but in DG. As in Figure 1, overlap was high during recall after fear conditioning (FC-Recall) and significantly reduced after extinction (EXT-Recall). Overlap remained low during the reinstating shock (IS), but increased during recall following the reinstating shock (IS-Recall) (n = 8-10 mice per group; F_3,31_ = 7.118, p = 0.0009; one-way ANOVA followed by pairwise comparisons corrected with the Benjamini-Hochberg method to correct for multiple comparisons; **P < 0.01, ns P > 0.05). **(h)** Quantification of relative numbers of eYFP+ cells across groups in the BLA (calculated as #eYFP+ cells divided by area in arbitrary units). No group had significantly different numbers of eYFP+ cells (n = 8-10 mice per group, F_3,32_ = 0.709, p = 0.554; one-way ANOVA). **(i)** same as **(h)** but in DG. No group had significantly different levels of eYFP+ cells (n = 8-10 mice per group, F_3,31_ = 2.404, p = 0.086; one-way ANOVA). **(j)** Quantification of relative numbers of cFos+ cells across groups in the BLA (calculated as #cFos+ cells divided by area in arbitrary units). Mice in the IS-Recall group had higher levels of cFos than mice in FC-Recall did (n = 8-10 mice per group, F_3,32_ = 3.926, p = 0.0171; one-way ANOVA followed by pairwise t-tests with Benjamini-Hochberg method to correct for multiple comparisons; *p < 0.05). **(k)** Same as **(j)** but in DG. No group had significantly different levels of cFos+ cells (n = 8-10 mice per group, F_3,31_ = 1.669, p = 0.194; one-way ANOVA).

**Supplementary Figure 2.**
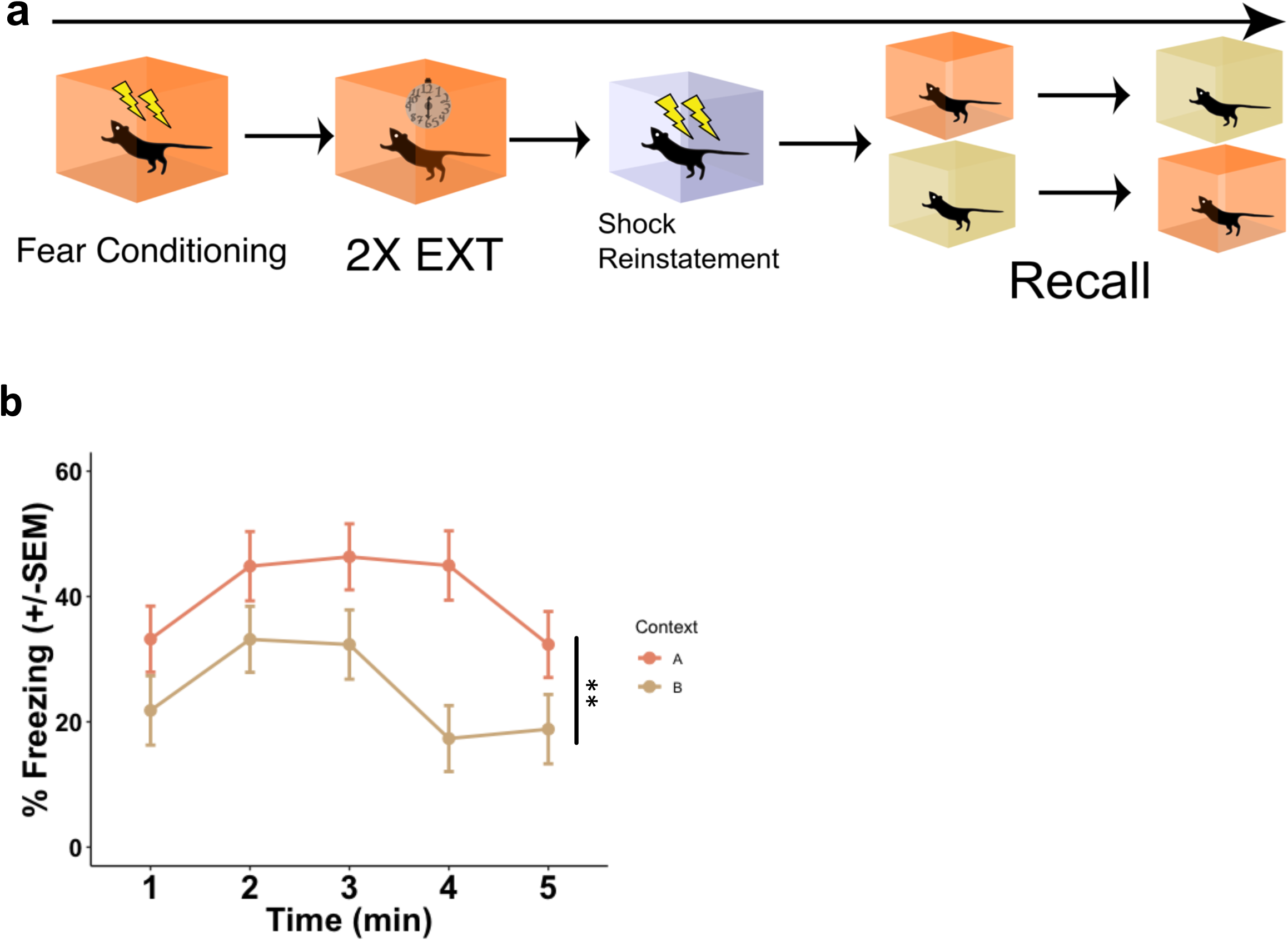
Reinstatement leads to partial generalization, but is largely context-specific. **(a)** Behavioral design. Mice underwent the reinstatement as described in Figure 1; however, they were placed in the original conditioned context (context A) and a novel context (context B), with some mice going into context A first and others context B first. **(b)** Compared to freezing in the novel context, mice froze significantly more in the original conditioned context across the 5-minute session (t_8_ = 3.415, **P = 0.0092; unpaired t-test; n = 5 minutes for each group).

**Supplementary Figure 3.**
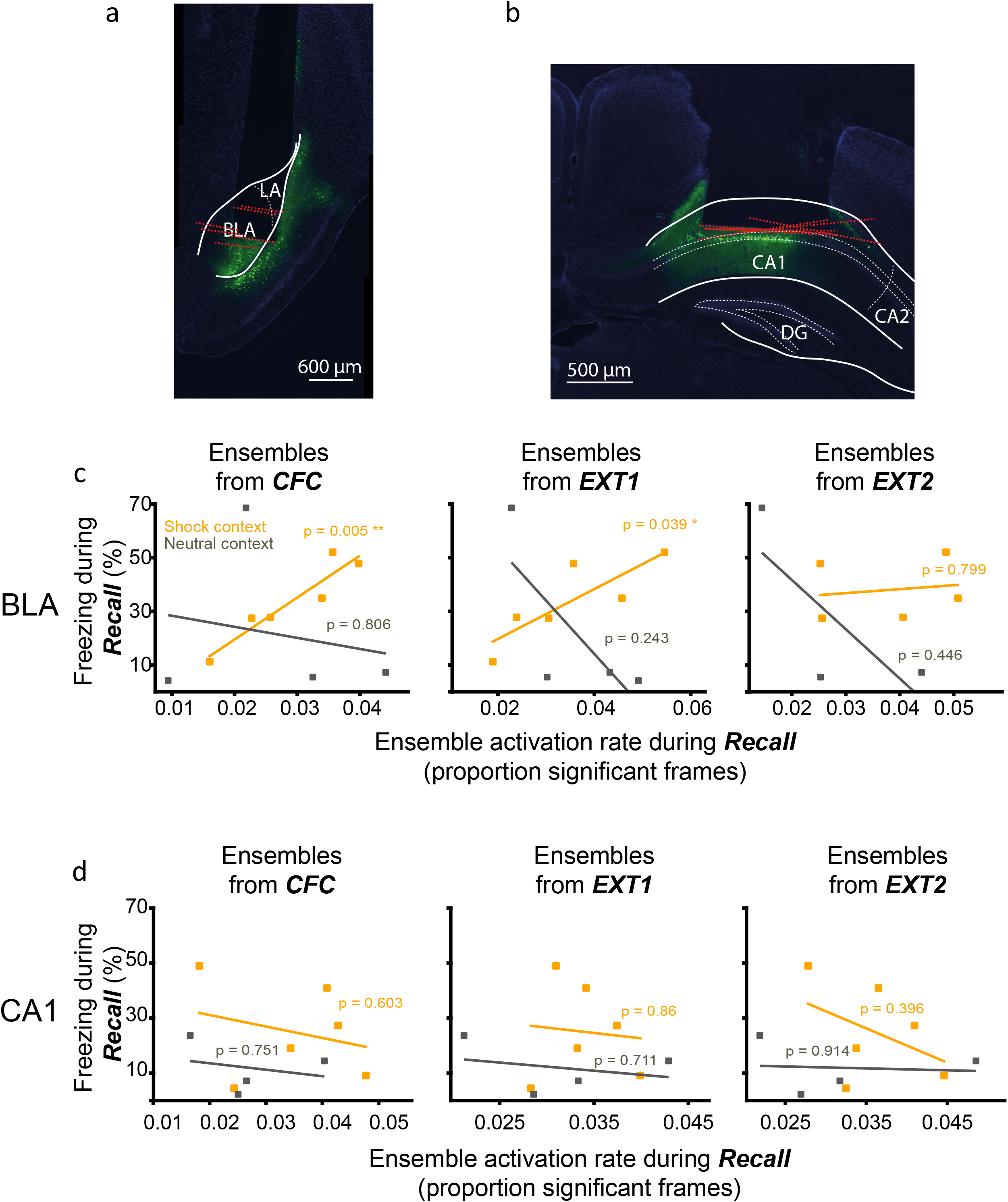
Reemergence of CFC-associated brain activity during fear relapse is specific to BLA and the shock context. **(a)** Reconstructed lens placements for all 6 BLA calcium imaging mice. Red dotted lines indicate the bottom of the lens. **(b)** Same as (**a**) but for the 6 CA1 mice. **(c)** Groupings of BLA neurons (ensembles) were extracted from CFC, EXT1, and EXT2 using a previously published PCA/ICA method (Lopes-dos-Santos et al., 2013). Averaged activations of those ensembles during Recall were correlated with freezing during Recall (CFC ensemble during Recall, Pearson correlation, P = 0.005; EXT1 ensemble during Recall, P = 0.039; EXT2 ensemble during Recall, P = 0.80). *n* = 6 mice (for the EXT2-Recall analysis, one mouse was omitted due to insufficient number of co-registered neurons). **(d)** Same as (**c**), but for CA1.

**Supplementary Figure 4.**
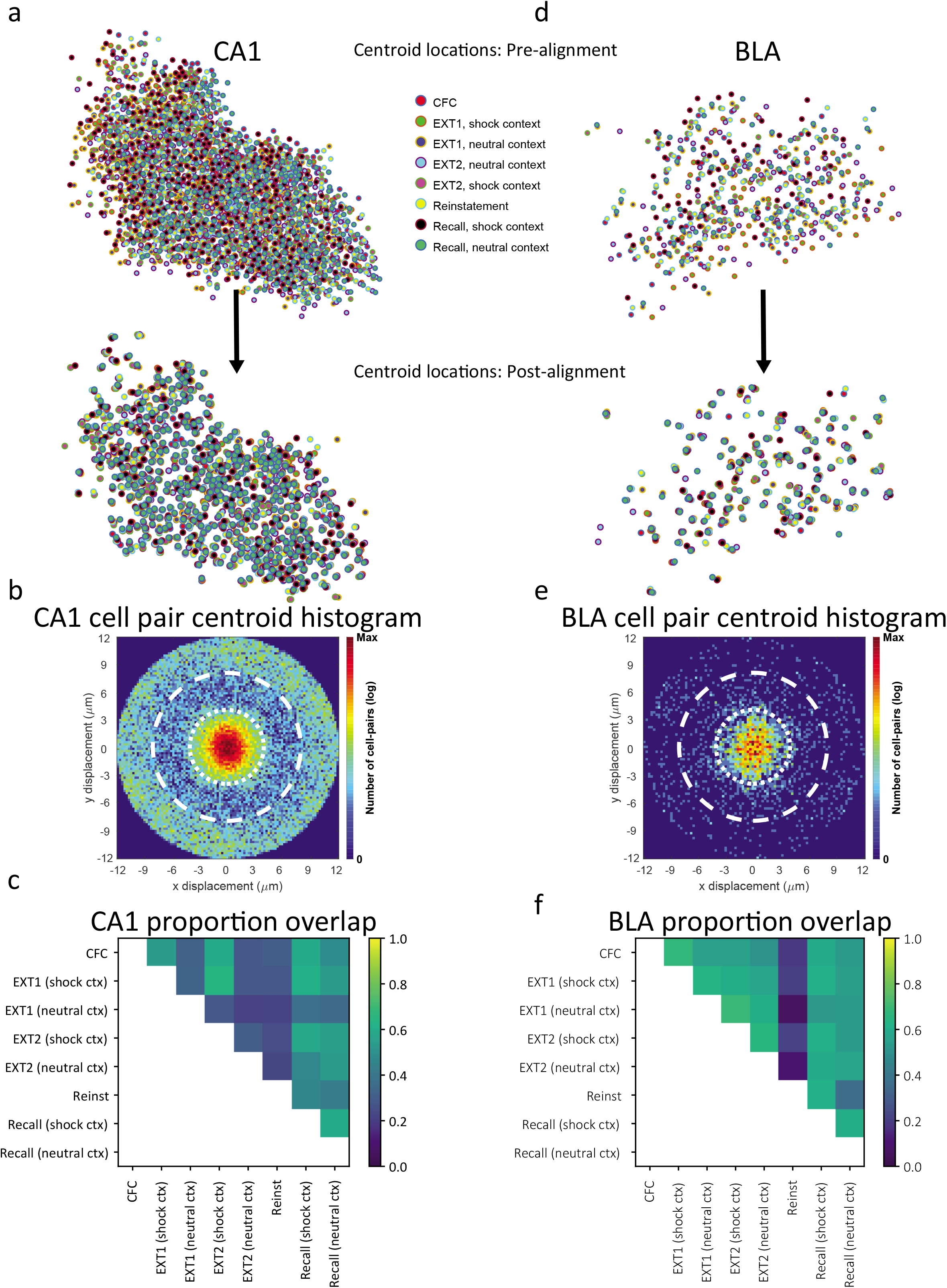
Cell registration and overlaps. **(a)** Registration diagnostics for example CA1 Ca^2+^ imaging mouse. Top, cell centroids prealignment. Bottom, cell centroids post-alignment. Colors correspond to different sessions. Figures were modified from plots generated via software from Sheintuch et al., (2017). **(b)** Two-dimensional histogram of matched pairs. Note the high density of cells within 3 microns of their match on another session. **(c)** Proportion of overlapping neurons (neurons with at least one calcium transient in the fear conditioning chamber in both sessions per session pair). There were no significant differences in overlaps across session pairs (all P > 0.06, Wilcoxon signed-rank test after corrections for multiple comparisons). **(d)** Same as (**a**), but for example BLA mouse. **(e)** Same as (**b**), but for example BLA mouse. **(f)** Same as (**c**), but for BLA mice.

**Supplementary Figure 5.**
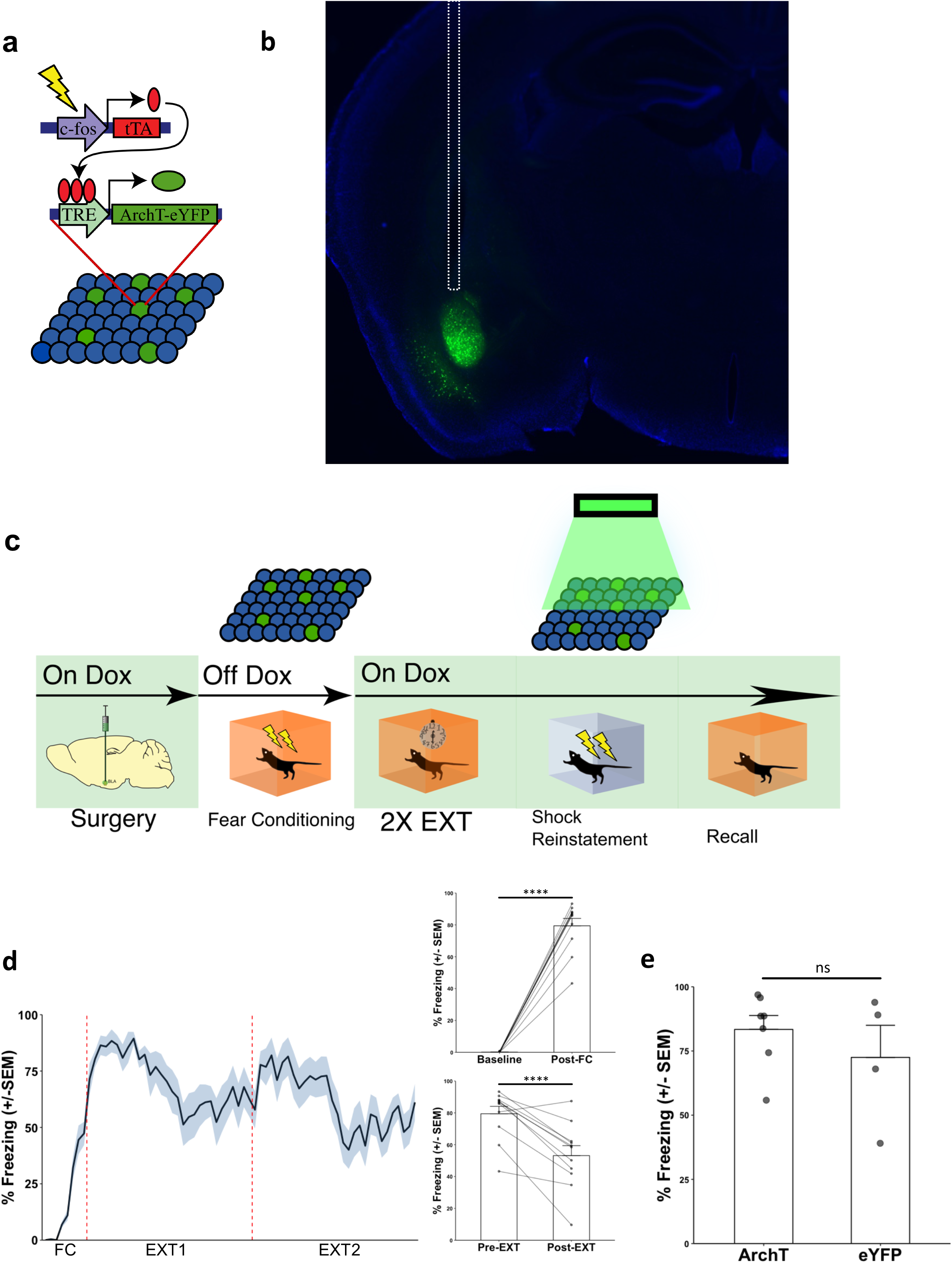
Inhibition of BLA fear ensemble does not prevent reinstatement. **(a)** Schematic of viral strategy. A viral cocktail of AAV9-c-Fos-tTA and AAV9-TRE-ArchT-eYFP was infused into the BLA for activity-dependent induction of ArchT-eYFP. Control mice received infusion of AAV9-c-Fos-tTA and AAV9-TRE-eYFP. **(b)** Representative microscope image of BLA injection site. Dotted line indicates optic fiber placement. **(c)** Behavioral schedule to test if inhibition of BLA fear ensemble during Shock Reinstatement can prevent reinstatement. **(d)** Left: Freezing during fear conditioning and extinction. Upper right: Compared to the first 3 minutes of FC (baseline freezing), mice froze significantly more in the first three minutes of the EXT1 session (t_10_ = 17.112, ****P < 0.0001; paired t-test; n = 10 mice). Bottom right: Mice spent significantly less time freezing in the last three minutes of EXT2 as compared to the first three minutes of EXT1 (t_10_ = 5.5159, ***P = 0.0003; paired t-test; n = 10 mice). **(e)** Compared to no-opsin controls (eYFP group), experimental mice that received optical inhibition (ArchT group) showed comparable levels of freezing during Recall, indicating that inhibition of the BLA fear ensemble did not prevent reinstatement (t_9_ = 0.935, n.s., P = 0.3742; unpaired t-test; ArchT, n = 7 mice; eYFP, n = 4 mice).

**Supplementary Figure 6.**
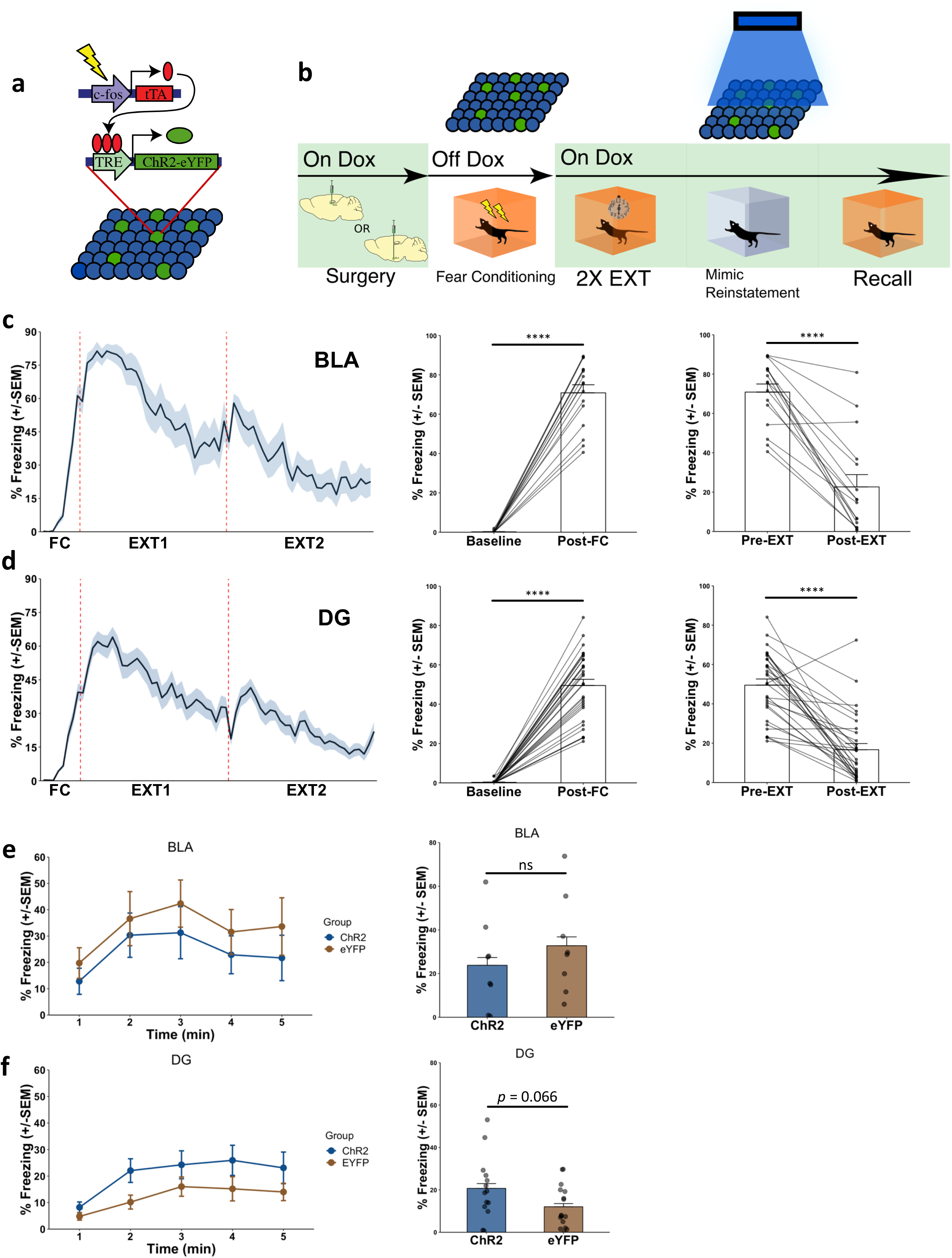
Stimulation of BLA or DG fear ensemble does not mimic reinstatement. **(a)** Schematic of viral strategy. A viral cocktail of AAV9-c-Fos-tTA and AAV9-TRE-ChR2-eYFP was infused into the BLA for activity-dependent induction of ArchT-eYFP. Control mice received infusion of AAV9-c-Fos-tTA and AAV9-TRE-eYFP. **(b)** Behavioral schedule to test if stimulation of BLA fear ensemble in a novel environment can mimic reinstatement. **(c)** Left: Freezing during fear conditioning and extinction for BLA-implanted animals. Middle: Compared to the first 3 minutes of FC (baseline freezing), mice froze significantly more in the first three minutes of the EXT1 session (t_15_= 17.171, ****P < 0.0001; paired t-test; n = 16 mice). Right: Mice spent significantly less time freezing in the last three minutes of EXT2 as compared to the first three minutes of EXT1 (t_15_= 7.9973, ****P < 0.0001; paired t-test; n = 16 mice). **(d)** Left: Freezing during fear conditioning and extinction for DG-implanted animals. Middle: Compared to the first 3 minutes of FC (baseline freezing), mice froze significantly more in the first three minutes of the EXT1 session (t_15_= 13.238, ****P < 0.0001; paired t-test; n = 16 mice). Right: Mice spent significantly less time freezing in the last three minutes of EXT2 as compared to the first three minutes of EXT1 (t_15_= 5.9671, ****P < 0.0001; paired t-test; n = 16 mice). **(e)** Left: Freezing across Recall session after BLA fear ensemble stimulation for ChR2 and eYFP groups. Right: Comparison of average freezing during Recall session after BLA fear ensemble stimulation, for ChR2 and eYFP groups (t_14_= 0.8265, n.s., P = 0.4224; unpaired t-test; n = 8 mice in each group). **(f)** Left: Freezing across Recall session after DG fear ensemble stimulation for ChR2 and eYFP groups. Comparison of average freezing during Recall session after DG fear ensemble stimulation, for ChR2 and eYFP groups (t_14_ = 1.9134, n.s., P = 0.06597; unpaired t-test; ChR2, n = 14 mice, eYFP, n = 16 mice).

**Supplementary Figure 7.**
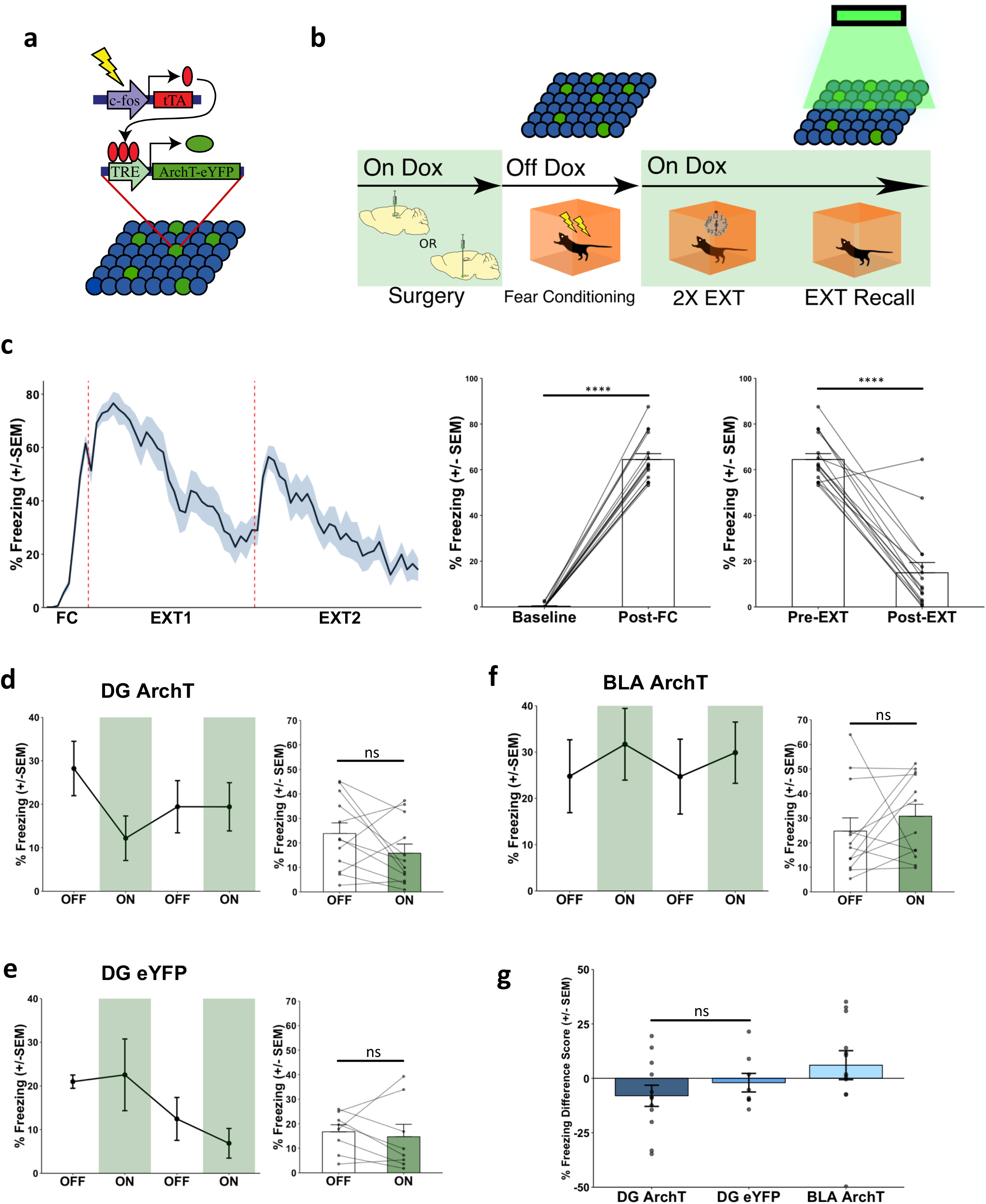
Inhibition of the fear ensemble after extinction does not alter freezing behavior. **(a)** Schematic of viral strategy. A viral cocktail of AAV9-c-Fos-tTA and AAV9-TRE-ArchT-eYFP was infused into the BLA for activity-dependent induction of ArchT-eYFP. Control mice received infusion of AAV9-c-Fos-tTA and AAV9-TRE-eYFP. **(b)** Behavioral design. Mice underwent FC and two EXT sessions, followed by an 8-minute Recall session, with 2-minute light OFF/light ON epochs. **(c)** Left: Freezing during fear conditioning and extinction. Middle: Compared to the first 3 minutes of FC (baseline freezing), mice froze significantly more in the first three minutes of the EXT1 session (t_15_= 25.898, ****P < 0.0001; paired t-test; n = 16 mice). Right: Mice spent significantly less time freezing in the last three minutes of EXT2 as compared to the first three minutes of EXT1 (t_15_= 9.7348, ****P < 0.0001; paired t-test; n = 16 mice). **(d-f)** Line graphs: 2-minute light OFF and ON epochs during Recall for the two experimental ArchT groups (DG ArchT & BLA ArchT) and the one control no-opsin group (DG eYFP). Bar graphs: Quantification of average freezing between light OFF vs. light ON epochs for each group. **(d)** DG ArchT Recall: mice froze no differently during ON epochs than OFF epochs (No main effect of Light: F_(1,5)_ = 2.306, n.s., P = 0.1893; two-way repeated measures ANOVA of Light (OFF vs ON) & Epoch (1 vs 2); n = 6 mice). **(e)** DG eYFP Recall: mice froze no differently during ON epochs than OFF epochs (No main effect of Light: F_(1,3)_ = 0.2404, n.s., P = 0.6575; two-way repeated measures ANOVA of Light (OFF vs ON) & Epoch (1 vs 2); n = 4 mice). **(f)** BLA ArchT Recall: mice froze no differently during ON epochs than OFF epochs (No main effect of Light: F_(1,5)_ = 0.6098, n.s., P = 0.4702; two-way repeated measures ANOVA of Light (OFF vs ON) & Epoch (1 vs 2); n = 6 mice). **(g)** Summary graph of freezing difference scores across all graphs in this figure, calculated as freezing in light ON epoch – freezing in light OFF epoch, for each set of epochs for each mouse. There was no significant difference in freezing between the DG ArchT and DG eYFP groups (t_18_ = 0.8689, n.s., P = 0.3963; unpaired t-test; DG ArchT, n = 12 scores from 6 mice; DG eYFP, n = 8 scores from 4 mice).

**Supplementary Table 1.** Cell counts in calcium imaging experiment.

| Animals | Sessions |
| --- | --- |

|  | CFC | EXT 1<br>(shock<br>context) | EXT 1<br>(neutral<br>context) | EXT 2<br>(shock<br>context) | EXT 2<br>(neutral<br>context) | Reinstatement | Recall<br>(shock<br>context) | Recall<br>(neutral<br>context) |
| --- | --- | --- | --- | --- | --- | --- | --- | --- |
| CA1 #1 | 858 | 851 | N/A | 858 | N/A | 849 | 845 | N/A |
| CA1 #2 | 734 | 754 | N/A | 745 | N/A | 727 | 727 | N/A |
| CA1 #3 | 714 | 735 | N/A | 736 | N/A | 685 | N/A | N/A |
| CA1 #4 | 464 | 435 | 432 | 457 | 405 | 405 | 422 | 386 |
| CA1 #5 | 597 | 558 | 399 | 367 | 311 | 531 | 650 | 664 |
| CA1 #6 | 517 | 454 | 417 | 492 | 441 | 468 | 513 | 555 |
| CA1 #7 | 473 | 419 | 292 | 321 | 302 | 440 | 390 | 466 |
| BLA #1 | 32 | 33 | N/A | 38 | N/A | 46 | 42 | N/A |
| BLA #2 | 84 | 75 | N/A | 91 | N/A | 86 | 86 | N/A |
| BLA #3 | 78 | 70 | 79 | 79 | 58 | 74 | 71 | 85 |
| BLA #4 | 52 | 59 | 37 | 36 | 32 | 39 | 39 | 44 |
| BLA #5 | 49 | 34 | 25 | 33 | 22 | 44 | 44 | 46 |
| BLA #6 | 19 | 10 | 9 | 16 | 4 | 10 | 27 | 22 |

## Methods and Materials

### Key Resources Table

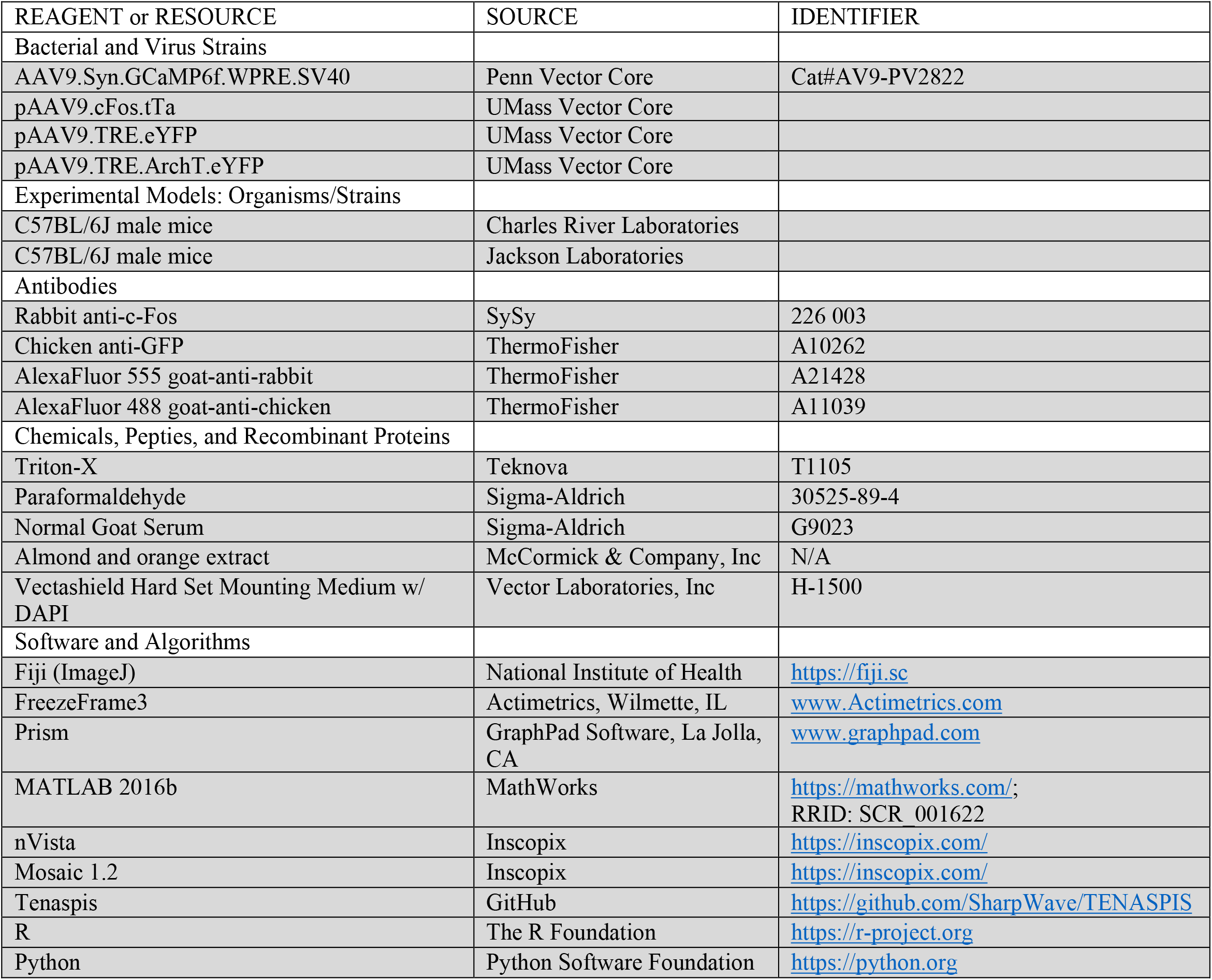

### Subjects

Wildtype male C57BL/6 mice (6-8 weeks of age; Charles River Labs) were housed in groups of 4-5 mice per cage. The animal facilities (vivarium and behavioral testing rooms) were maintained on a 12:12-hour light cycle (lights on at 0700). Mice were placed on a diet containing 40 mg/kg doxycycline (Dox) for a minimum of two days before receiving surgery with access to food and water *ad libitum*. Mice recovered for at least ten days after surgery. Dox-containing diet was replaced with standard mouse chow (*ad libitum*) 48 hours prior to behavioral tagging to open a time window of activity-dependent labeling (Ramirez et al., 2013).

All procedures relating to mouse care and treatment conformed to the institutional and National Institutes of Health guidelines for the Care and Use of Laboratory Animals. No statistical methods were used to predetermine sample size; however, sample sizes were chosen based on sample sizes in previous studies (Ramirez et al., 2013). Data collection and analysis were not performed blind to the conditions of the experiments.

### Activity-dependent viral constructs

pAAV_9_-cFos-tTA (titer of ∼1.5 × 10^13^ GC/mL), pAAV_9_-TRE-eYFP (titer of ∼1 × 10^13^ GC/mL), and pAAV 9 -TRE-ArchT-eYFP (titer of ∼1 × 10^13^ GC/mL) were constructed as previously described (Ramirez et al., 2015). pAAV_9_-c-Fos-tTA was combined with pAAV_9_-TRE-eYFP or pAAV_9_-TRE-ArchT-eYFP prior to injection at a 1:1 ratio.

### Stereotaxic surgeries

#### Opsin injections and optic fiber implants

Stereotaxic injections and optical fiber implants followed methods previously reported(Ramirez et al., 2015). All surgeries were performed under stereotaxic guidance and subsequent coordinates are given relative to Bregma (in mm). Mice were mounted into a stereotaxic frame (Kopf Instruments, Tujunga, CA, USA) and anesthetized with 3% isoflurane during induction and lowered to 1-2% to maintain anesthesia (oxygen 1L/min) throughout the surgery. Ophthalmic ointment was applied to both eyes to prevent corneal desiccation. Hair was removed with scissors and the surgical site was cleaned with ethanol and betadine. Following this, an incision was made to expose the skull. Bilateral craniotomies involved drilling windows through the skull above the injection sites using a 0.5 mm diameter drill bit. Coordinates were -1.35mm anteroposterior (AP), ±3.45mm mediolateral (ML), and -5.15mm dorsoventral (DV) for basolateral amygdala (BLA)(Davis et al., 2017); -2.2mm AP, ±1.3mm ML, and -2.0mm DV for dorsal dentate gyrus (dDG)(Ramirez et al., 2015); and -2.0mm AP, ±1.5mm ML, and -1.55mm DV for dorsal CA1 (dCA1). All mice were injected with a volume of 0.3 μl of AAV9 cocktail per site at a control rate of 0.1 μl min^-1^ using a mineral oil-filled 33-gage beveled needle attached to a 10 μl Hamilton microsyringe (701LT; Hamilton) in a microsyringe pump (UMP3; WPI). The needle remained at the target site for two minutes post-injection before removal. For dDG optogenetic experiments, a bilateral optic fiber implant (200 μm core diameter; Doric Lenses) was chronically implanted above the injection site (−1.6mm DV). For BLA optogenetic experiments, monofibers were implanted above each injection site (−4.9mm DV). For CA1 optogenetic experiments, a bilateral optic fiber implant (200 μm core diameter; Doric Lenses) was chronically implanted above the injection site (−1.15mm DV). Jewelry screws secured to the skull acted as anchors. Layers of adhesive cement (C&B Metabond) followed by dental cement (A-M Systems) were spread over the surgical site. Mice that did not receive implants had their incision sutured. Mice received 0.1 mL of 0.3 mg/ml buprenorphine (intraperitoneally) following surgery and were placed on a heating pad during recovery.

#### GCaMP6f injections and lens implants

Mice in Ca^2+^ imaging experiments underwent three separate serial surgeries. First, mice received unilateral infusions of AAV9-Syn-GCaMP6f (U Penn Vector Core, titer of ∼5 × 10^12^ GC/mL) into either right CA1 (AP -2.0 mm, ML +1.5 mm, DV -1.5 mm) or right BLA (AP -1.35 mm, ML +3.45 mm, DV -5.05 mm). The viral vector was injected at a rate of 40 nL/min and allowed 10 min to diffuse before the scalp was sutured.

Two to four weeks after viral infusion, mice were implanted with a gradient index (GRIN) lens into either CA1 (1 mm diameter, 4 mm length, Inscopix; AP -2.25 mm, ML +1.8 mm, DV -1.3 mm) or BLA (0.65 mm diameter, 7.3 mm length; AP -1.25 mm, ML +3.15 mm, DV -4.85 mm). For CA1 implants, overlying neocortex was aspirated under continuous irrigation with cold 0.9% saline until vertical white fibers were visible (Resendez et al., 2016). For BLA implants, a tract was created using a stereotaxically lowered 27-gauge needle (0.5 mm diameter) into the craniotomy prior to insertion of the lens. Gaps between the lens and the skull were filled using Kwik-Sil (World Precision Instruments) and the lens was then adhered to the skull using Metabond. The surface of the lens was covered with a protective cap made of Kwik-Cast (World Precision Instruments) until base plate attachment.

Finally, one week after the lens implant, mice were implanted with a base plate for microscope attachment. A plastic base plate was magnetically attached to the bottom of the microscope. The microscope objective was then aligned to the GRIN lens and lowered until cells came into focus, as observed via nVista recording software (Inscopix). The base plate was then adhered to the surrounding Metabond on the animal’s skull using a dental composite (Flow-It ALC, Pentron) and strengthened with an additional layer of Metabond.

Histological assessment verified viral targeting and fiber/lens placement. Data from off-target injections and implants were not included in analyses.

### Optogenetic methods

Optic fiber implants were plugged into a patch cord connected to a 520nm green laser diode controlled by automated software (Doric Lenses). Laser diode output was tested at the beginning of every experiment to ensure that at least 10 mW of power was delivered at the end of the optic fiber tip (Doric lenses). Mice began the stimulation trial with a 2-min light-off epoch, followed by 2-min optical stimulation (15 ms pulse width, 20 Hz), and then repeated, such that the mice underwent a light-OFF/ON/OFF/ON pattern for a total of 8-min.

### Behavioral tagging

Dox diet was replaced with standard lab chow (*ad libitum*) 48-hours prior to behavioral tagging. *Female exposure*: One female mouse (PD 30-40) was placed into a clean home cage with a clear cage top, which was used as the interaction chamber. The experimental male mouse was then placed into the chamber and allowed to interact freely for one hour(Ramirez et al., 2015). *Fear conditioning*: Mice were placed into a conditioning chamber and received fear conditioning (see below) over a 500-second training session (including exposure to four 1.5 mA foot shocks). Following behavioral tagging, the male mouse was returned to their home cage with access to Dox diet (Ramirez et al., 2013).

### Behavior

All behavior assays were conducted during the light cycle of the day (0730–1930). Mice were handled for 1-2 days, 2 min per day, before all behavioral experiments, and were run by cage. The entire behavioral schedule includes female exposure, fear conditioning, extinction, reinstatement, and recall (described below). Which of these behaviors the mice underwent depended on the experiment.

#### Female exposure

One female mouse (PD 30-40) was placed into a clean home cage with a clear cage top and no bedding, which was used as the interaction chamber. The experimental male mouse was then placed into the chamber and allowed to interact freely for one hour(Ramirez et al., 2015).

#### Fear conditioning

Fear conditioning occurred in one of four mouse conditioning chambers (Coulbourn Instruments, Whitehall, PA, USA) with metal-panel side walls, Plexiglas front and rear walls, and a stainless-steel grid floor composed of 16 grid bars. The grid floor was connected to a precision animal shocker (Coulbourn Instruments, Whitehall, PA, USA) set to deliver a 2-second 1.5 mA foot shock unconditioned stimulus (US). A ceiling-mounted video camera recorded activity and fed into a computer running FreezeFrame3 software (Actimetrics, Wilmette, IL, USA). The software controlled stimuli presentations and recorded videos from four chambers simultaneously. The program determined movement as changes in pixel luminance. Context alterations included changes to spatial, olfactory, tactile, and lighting cues. The conditioning chamber with room lights off was designated as Context A. Context B involved modifications to the conditioning chamber, including vertical black and white strips spaced ∼ 3 cm apart obscuring the front and rear walls, black inserts placed between grids to slightly alter dimensions of the box, 1 mL of almond extract in a plastic container positioned below the grid floor, and room lights on. Context C also involved modifications to the conditioning chamber, with a plastic sheet with a cross-hatch texture placed over the shock grid to change tactile cues, a black sheet obscuring the front walls, 1 mL of orange extract in a plastic container position below the grid floor, and room lights on. The chambers were cleaned with 70% ethanol solution prior to animal placement. Contextual fear conditioning occurred in Context A. Briefly, mice were placed into the conditioning context for a 500-second acquisition session, including a 180- second baseline period followed four 1.5 mA, 2-second foot shock USs (interstimulus interval [ISI] equals 80-sec). In optogenetic experiments, mice had patch cords attached near the conditioning chamber by the experimenter, and were run two mice at a time.

Fear conditioning data are collected using FreezeFrame3 software (Actimetrics, Wilmette IL) with the bout length set at 1.25-sec and the freezing threshold initially set as described in the program instructions. Freezing is defined as changes in pixel luminance falling below a threshold. An experimenter adjusted the threshold so that freezing behavior involves the absence of all movement except those needed for respiration as previously described. Freezing behavior was scored as the percentage of time spent freezing during a given bout of time. Statistical analyses involved paired t-tests comparing within subject differences (i.e. light off vs light on epochs), unpaired t-tests comparing across experimental groups (e.g. ArchT group vs. eYFP group), and one-sample t-tests comparing freezing differences scores to a µ_0_ = 0.

#### Extinction

Extinction occurred in Context A (described above) the day following fear conditioning. Mice were placed in Context A for 30-min sessions once per day, for two days. As in fear conditioning, cages of four mice were run simultaneously, and cages of five mice were run as three mice first, then the remaining two.

#### Reinstatement

Reinstatement occurred in Context B (described above) the day following the second day of extinction. Mice were placed in Context B and given a 1.5 mA, 2-second foot shock 1-second into the trial. Mice were left in the chamber for another 60-seconds before being removed. As opposed to running four mice simultaneously as in fear conditioning and extinction, each mouse in a cage of 4 mice was run individually for reinstatement.

#### Recall

Recall for behavioral and overlap experiments involved placement in a context for 5-min. In this case, as in fear conditioning and extinction, cages of four mice were run simultaneously while cages of five mice were run as three mice first, then the remaining two. In optogenetic experiments, recall involved an 8-min session consisting of 2-min epochs of alternating light off and light on. In this case, mice were run one or two at a time.

### Immunohistochemistry

Mice were anesthetized with 3% isoflurane and perfused transcardially with cold (4° C) phosphate-buffered saline (PBS) followed by cold 4% paraformaldehyde (PFA) in PBS. Brains were extracted and stored overnight in PFA at 4°C. Fifty μm coronal sections were collected in serial order using a vibratome and collected in cold PBS (100 μm coronal sections were collected when solely verifying injection site and implant placement). Immunostaining involved washing sections in PBS with 0.2% triton (PBST) for 10-minutes (x3). Sections were blocked for 1 hour at room temperature in PBST and 5% normal goat serum (NGS) on a shaker. Sections were transferred to wells containing primary antibodies (1:5000 rabbit anti-c-Fos [SySy]; 1:500 chicken anti-GFP [Invitrogen]) and allowed to incubate on a shaker overnight at 4°C. Sections were then washed in PBST for 10-min (x3), followed by 2-hour incubation with secondary antibody (1:200 Alexa 555 anti-rabbit [Invitrogen]; 1:200 Alexa 488 anti-chicken [Invitrogen]). Following three additional 10-min washes in PBST, sections were mounted onto micro slides (VWR International, LLC). Vectashield Hard Set Mounting Medium with DAPI (Vector Laboratories, Inc) was applied, slides were cover slipped, and allowed to dry overnight.

### Cell counting

The number of eYFP- or cFos-immunoreactive neurons in the dentate gyrus and basolateral amygdala were counted to measure the number of active cells during defined behavioral tasks per mouse. Only animals that had accurate injections were selected for counting, and sections closest to the injection coordinates for each brain region were chosen (see the Stereotaxic surgeries section of Methods). All animals were sacrificed 90 minutes post-assay for immunohistochemical analyses. Fluorescence images were acquired using a confocal microscope (Zeiss LSM800, Germany) with a 10X objective for dentate gyrus images and 20X objective for basolateral amygdala images. Z-stacked images were taken at step sizes of 3um for DG and 1.54um for BLA in the z-axis and maximum projections of these z-stacks were taken for counting.

For the dentate gyrus, the number of eYFP-positive and cFos-positive cells in a set region of interest were quantified manually using ImageJ. eYFP positive cells in the basolateral amygdala were quantified automatically using a custom Python script that detects cells based on size and circularity of objects in an image, while cFos-positive cells were quantified manually with ImageJ because of poorer signal to noise in cFos images compared with the eYFP images.

To calculate the percentage of re-activated cells we counted the number of eYFP-positive cells, cFos-positive cells, and both eYFP- and cFos-positive (Overlapped) cells. Re-activation was calculated as (Overlap/eYFP*100). Overlap was compared across groups using unpaired t-test (two-groups) and one-way ANOVA (more than two groups). Relative expression of eYFP and cFos across groups was calculated as (#eYFP cells/Area) and (#cFos cell/Area) in arbitrary area units.

### *In vivo* calcium imaging

A miniaturized microscope (Inscopix) was used to collect Ca^2+^ imaging videos in mice undergoing the fear reinstatement schedule. Videos were captured using nVista (Inscopix) at 20 Hz in a 720 x 540 pixel field of view (1.1 microns/pixel). Microscope attachments were done while the mice were awake and restrained. Mice experienced both conditioned (A) and neutral (B) context exposures during extinction and reinstatement recall in a counterbalanced fashion; that is, if a mouse experienced context A first on extinction day 1, they experienced context B first on extinction day 2 and context A first again on reinstatement recall. Other mice experienced context B first on extinction day 1, and so on.

Ca^2+^ imaging videos were cropped, spatially bandpass filtered, and motion corrected offline using Inscopix Data Processing Software v1.1. A ΔF/F movie was computed using the mean fluorescence of the movie as the baseline and PCA/ICA was used for automated segmentation of cell masks (Mukamel, Nimmerjahn, & Schnitzer, 2009). PCA/ICA putative cell masks were each manually inspected to verify that cells were accurately captured with high fidelity. Cells across imaging sessions were aligned and registered to each other using the automated CellReg software in Matlab (Sheintuch et al., 2017).

Population vectors (PVs) were computed for the entirety of the CFC session by taking the average Ca^2+^ transient rate for each cell while the mouse was in the fear conditioning chamber. PVs for EXT1, EXT2, and Recall were defined as the average Ca^2+^ transient rate for each 30 s time bin while the mouse was in the chamber. As a measure of the similarity of the population to the CFC network state, Pearson correlations were performed between each 30 s PV to the CFC PV. As a control, we also performed correlations between PVs from a neutral context to the CFC PV. Of the 6 BLA mice and 6 CA1 mice, 4 mice from each group were given these neutral context exposures. Only cells that were active for both sessions being correlated (CFC and either EXT1, EXT2, or Recall) were considered.

We used regression discontinuity analysis to statistically detect deflections in the PV correlation trajectory across time using the rdd package available for Python (https://pypi.org/project/rdd/). We defined two possible boundaries that could potentially yield a statistically significant effect – the time points separating EXT1/EXT2, and those separating EXT2/Recall. Next, we fit an ordinary least squares model with time and each boundary as predictors. A discontinuity was inferred if the p-value of the boundary predictor was < 0.05, Bonferroni-corrected for multiple comparisons.

We used piecewise regressions to determine whether reinstatement increased PV correlations (to CFC) greater than would be expected from chance. For each mouse, we fit PV correlation coefficients using time as a predictor for the EXT sessions and the Recall session separately. Next, using the regression models obtained from the EXT session data, we predicted the PV correlation value that should be observed during the first time bin in Recall. Then we compared the predicted values to the actual values derived from the y-intercepts of the regression models on the Recall data.

### Neuronal Ensembles

We characterized neuronal ensembles from Ca^2+^ transient activity and correlated their later activity during Recall to relapse-associated freezing. To extract neuronal ensembles, we used a PCA/ICA method that was previously published (Lopes-dos-Santos et al., 2013). This method is a source separation algorithm that is similar in principle to the PCA/ICA that was run on Ca^2+^ imaging movies earlier (Mukamel et al., 2009). However, rather than extracting neurons from pixel fluorescence, we are now extracting ensembles from traces. First, Ca^2+^ traces were z-score normalized. Then, principal components analysis (PCA) was used to reduce the dimensionality of the trace matrix and to compute eigenvalues corresponding to the variances along the reduced dimensions. To produce a surrogate dataset for determining significant eigenvalues, we circularly shuffled each neuron’s Ca^2+^ trace vector and performed PCA on that shuffled data 1000 times. Next, significant ensembles were tallied based on eigenvalues exceeding a statistically defined threshold (eigenvalues must be > 99% of all the eigenvalues computed from the PCA of the shuffled data). To extract statistically independent ensembles, an independent components analysis (ICA) was run on the trace matrix with the number of components specified by the number of statistically significant ensembles (eigenvalues). From this computation, each ensemble now has a corresponding “pattern” vector describing the weight of each neuron contributing to that ensemble. For each ensemble, a projection matrix was computed from the outer product of these pattern vectors with the identity of the matrix set to 0 (to ensure that more than one neuron was needed to trigger an ensemble activation). The projection matrix was multiplied with each observation (frame) of the trace matrix to obtain ensemble activation strength. Finally, an ensemble was considered to be significantly activated if it exceeded 2 standard deviations above its mean activation strength.

Ensembles were extracted individually for CFC, EXT1, and EXT2. Only neurons that were active between these sessions and Recall were considered. After ensembles were extracted, their activity strength during Recall was calculated and the proportion of significant activations was correlated with freezing.

### Data Analysis

Data were analyzed using Inscopix nVista in conjunction with custom-made R, Python, and Matlab scripts. Data were analyzed using paired t-tests (two factors) or with one-way and repeated measures ANOVAs (more than two factors), Wilcoxon signed-rank tests, and Mann-Whitney U tests (two-tailed, corrected for multiple comparisons using false discovery rate adjustments). Post-hoc analyses (Tukey’s) were used to characterize treatment and interaction effects, when statistically significant (alpha set at p<0.05, two-tailed).

